# A comprehensive genetic study of vaginal microbiota composition of Holstein cows

**DOI:** 10.64898/2026.10.09.757891

**Authors:** Abdullah Al Momen Sabuj, Pascal Croiseau, Sébastien Ducrocq, Sophie Merlin, Sophie Martel, Christophe Audebert, Jordi Estellé, Marie-Pierre Sanchez, Louise Brulin

## Abstract

The vaginal microbiota is a potentially important component of the bovine reproductive tract and has been associated with fertility, health, and production-related traits in cattle. However, the impact of host genetics on the bovine vaginal microbiota and its genetic correlations with animal performance remain largely unexplored. To investigate this, we performed a comprehensive genetic study of the vaginal microbiota in Holstein cows. Vaginal swabs from 681 lactating cows across 18 herds in Northern France were collected, along with records of fertility, milk production, and health traits. The microbiota samples were sequenced for the 16S rRNA gene, generating count tables of amplicon sequence variants (ASV).

We estimated genetic parameters for vaginal microbiota α-diversity (Chao1, Shannon, and Simpson indices) and β-diversity (Bray-Curtis dissimilarity matrix), as well as the centered log-ratio (CLR)-transformed abundances of taxa present in more than 50% of samples. Across the 173 analyzed taxa, heritability estimates ranged from low to moderate (h² = 0.09 to 0.38) for five genera, three families, eight orders, three classes, and one phylum. Notably, the order Flavobacteriales exhibited the highest heritability estimate. The study also highlighted genetic correlations between heritable microbiota traits and host phenotypes related to fertility, production, and health. Sequence-based genome-wide association studies (GWAS) identified biologically relevant signals, including a quantitative trait locus (QTL) for the genus *Alloprevotella* on chromosome 6, with lead variants located within the *NAF1* gene. Additionally, two signals on chromosome 17 for the Bacteroidales order and the Bacteroida class co-localized with *IL15*.

Our findings show that host genetics contributes to variation in specific components of the bovine vaginal microbiota and suggest that microbiome-related traits may warrant further investigation as potential indicators in dairy cattle breeding and management research.

## Introduction

Microbiota comprises a diverse community of microorganisms residing in specific environments, including bacteria, archaea, fungi, viruses, and protozoans. These microbial communities are found in various locations of animals, humans, and plants, significantly influencing host health, development, physiology, and behavior (Parfrey et al., 2018; Appiah et al., 2020). In ruminants, microbiota are found in multiple tissues, including the skin, oral cavity, rumen, lower gut, and reproductive tract, each playing a crucial role in the host’s overall health and well-being (Li et al., 2019; Barden et al., 2020; Bay et al., 2023).

In bovine research, particular attention has been directed toward understanding the composition and functions of the gut microbiota, especially in the rumen and lower gastrointestinal tract, as they play pivotal roles in maintaining animal health and optimizing production (O’Hara et al., 2020; Amat et al., 2022). Recently, the microbiota of the reproductive tract has received increased attention, as reproductive success is critical to the profitability of dairy and beef cattle (Ault-Seay et al., 2023; Zangirolamo et al., 2024). Moreover, focusing solely on production traits, such as milk and meat, can often lead to cows with lower reproductive efficiency and poorer health (Weigel et al., 2017; Cardoso Consentini et al., 2021). Studies have indicated that the estimated annual losses from reproductive failure amount to 100€ per cow (van Soest et al., 2019; Wicaksono et al., 2025). While reproductive technologies and management strategies involving genetic selection and nutrition positively impact reproductive efficiency in dairy cattle (Giordano et al., 2011; Cardoso Consentini et al., 2021), insights into the reproductive microbiota also play a role in understanding reproductive success and failure (Adnane and Chapwanya, 2022; Brulin et al., 2024b; Gupta et al., 2024).

The bovine vagina harbors a diverse and unique microbial community distinct from that of the upper reproductive tract (Rodrigues et al., 2015; Amat et al., 2022). Its composition appears to be influenced by host and reproductive factors, including parity and physiological status (Ni et al., 2023; Dias et al., 2024). Microorganisms help maintain the acidic pH of the vagina, thereby preventing the growth of harmful bacteria and promoting the growth of beneficial ones (Miller et al., 2016; Kovachev, 2018; Punzón-Jiménez and Labarta, 2021).

Among dairy cattle, the predominant phyla in the vaginal microbiota include Firmicutes, Bacteroidetes, Proteobacteria, and Actinobacteria (Deng et al., 2019; Chen et al., 2020; Brulin et al., 2024b). Additionally, some studies have reported that Tenericutes, Euryarchaeota, and Fusobacteria are abundant (Chen et al., 2020; Quadros et al., 2020; Moreno et al., 2022). Genera such as *Ureaplassma, Mycoplasma, Bacteroidales*, and *Clostridium* are frequently observed in the vaginal microbiome of cattle and have been linked to reproductive disorders (Deng et al., 2019; Kudo et al., 2021). Certain bacteria commonly found in the gut, such as *Bacteroides*, *Ruminococcus*, *Clostridium*, *Enterobacteriaceae*, and *Prevotella*, are also present in the vaginal microbiota, potentially due to the proximity of the vagina and anus and to frequent contact between feces and the vulva (Laguardia-Nascimento et al., 2015). Unlike in humans, where the vaginal microbiota is primarily dominated by *Lactobacillus spp*. *Lactobacillus* is less commonly identified and typically present at low abundance (<1%) in dairy cattle (Sirota et al., 2014; Swartz et al., 2014; Clemmons et al., 2017). This marked difference between bovine and human vaginal microbiota highlights the importance of species-specific studies. In recent research, we have revealed significant associations between the abundance of cow vaginal microbiota and host traits related to health, dairy production, and fertility (Brulin et al., 2024b).

Host genetics play a crucial role in shaping the microbial communities associated with the host, including those inhabiting the vagina. Numerous studies have demonstrated low to moderate heritability in vaginal microbiota in pigs (Sanglard et al., 2020) and humans (Si et al., 2017; Mccauley et al., 2022). Genome-Wide Association Studies (GWAS) have further identified links between specific genomic regions and the vaginal microbiota (Sanglard et al., 2020; Wright et al., 2021). However, there is a shortage of information regarding the heritability and genotype associations of vaginal microbiota in cows. Addressing this knowledge gap is crucial to understanding how host genetics influences the vaginal microbiota and to exploring strategies to manage it, thereby enhancing reproductive success and health in dairy cattle.

In this study, we hypothesized that host genetics influences the composition of the vaginal microbiota in Holstein cows and that genetic correlations exist between this bacterial community and host phenotypes. Furthermore, we hypothesized that specific regions in the cow genome could contribute to variation in vaginal microbial composition. To test these hypotheses, we aimed to estimate the heritability of vaginal microbiota composition, i.e., the proportion of variation controlled by host genetics. We also sought to estimate genetic correlations between microbial taxa and host traits, including fertility, production, and health. Additionally, we aimed to identify specific genes and genomic variants that contribute to the genetic determinants of the vaginal microbiota through GWAS of imputed whole-genome sequence data.

## Materials and Methods

### Animal Sampling and Metadata

A total of 681 vaginal swab samples were collected from Holstein cows in 18 commercial herds in Northern France between September 2017 and December 2018, following the procedure described in Brulin et al. (2024b). Briefly, artificial insemination (AI) technicians affiliated with Genes Diffusion (Douai, France) collected samples between calving and pre-conception, and all samples were stored at −20°C in sterile tubes at Genes Diffusion’s research facilities (Institut Pasteur de Lille, France) until DNA extraction.

Across the 18 herds, an average of 38 ± 17 samples per herd was obtained, ranging from 17 to a maximum of 84. Samples were collected from cows with parity ranging from 1 to 5. The highest number of samples came from parity 1 (236), while the lowest was from parity 5 (30). The period from 20 to 425 days post-calving was divided into nine categories, each spanning a 20-day interval. On average, 76 samples were obtained per category, as summarized in Table 1. Samples were collected during three distinct seasons (winter, spring, autumn).

**Table 1:** Descriptive statistics of different variables.

| Variables | Category | Average animals | SD | Max | Min |
| --- | --- | --- | --- | --- | --- |
| Herd | 18 | 38 | 17 | 84 | 17 |
| Parity | 5 | 136 | 94 | 263 | 30 |
| Sampling season | 3 | 227 | 107 | 340 | 136 |
| Sequencing run | 4 | 170 | 60 | 229 | 94 |
| Calving to sampling interval | 9 | 76 | 59 | 204 | 25 |
| Adjusted Host Phenotypes | Samples | Mean | SD | Max | Min |
| Calving-first AI interval (CAIf) | 666 | 4.91 | 21.68 | 98.66 | -54.19 |
| First AI to last AI (FLAI) | 664 | 7.56 | 45.79 | 301.88 | -69.34 |
| Cow conception rate (CCR) | 666 | 0.02 | 0.24 | 0.58 | -0.58 |
| Milk yield (MY) | 668 | -118.49 | 679.37 | 1829.24 | -4910.84 |
| Protein yield (PY) | 668 | -7.21 | 19.83 | 45.27 | -156.48 |
| Fat yield (FY) | 668 | -10.08 | 28.71 | 69.59 | -197.02 |
| Protein content (PC) | 668 | -0.51 | 1.65 | 5.01 | -5.91 |
| Fat content (FC) | 668 | -0.76 | 3.42 | 10.18 | -11.22 |
| Clinical mastitis (CMA) | 512 | 0.01 | 0.25 | 1.10 | -0.49 |
| Somatic cell count score (SCC) | 668 | 0.21 | 0.81 | 6.33 | -1.48 |

\*SD= Standard deviation; Max=Maximum; Min=Minimum

Phenotypic data on cows’ fertility, productivity, and health were recorded at the time of sample collection and adjusted for non-genetic effects as previously described in Brulin et al. 2024. Fertility parameters considered were: calving-to-fertilization interval via AI (CAIf), interval between the first and last AI (FLAI), and cow conception rate (CCR). Furthermore, various production traits were documented, including milk yield (MY), protein yield (PY), and fat yield (FY), along with their associated contents, protein content (PC) and fat content (FC), measured over the initial 305 days of lactation. For mammary health assessment, clinical mastitis (CMA) and somatic cell count (SCC) were considered. To account for environmental effects, phenotypes were adjusted for non-genetic effects and expressed as yield deviations (YD). The French national genetic evaluation system calculated these YDs for the entire Holstein population using the models described at https://interbull.org/ib/geforms. A summary of traits is presented in Table 1.

### DNA Extraction, 16S rRNA Gene Sequencing, and ASV Processing

For DNA extraction and sequencing, samples were divided into four laboratory batches, each containing 94-229 samples. DNA extraction, library preparation, and high-throughput sequencing were performed as previously described by Brulin et al. (2024b). Briefly, DNA was extracted using a commercial extraction kit. Libraries were prepared using a dual-indexed paired-end sequencing approach targeting the V3-V4 variable regions of the 16S rRNA gene and sequenced on the Illumina MiSeq platform with the MiSeq Reagent Kit v3 (600 cycles). Sequence quality was assessed with FastQC (v.0.11.9).

Sequence analyses were performed using the dada2 (v1.24-0) pipeline in R as outlined by Brulin et al. (2024b). Reads were filtered, denoised, merged, and checked for chimeras before defining Amplicon Sequence Variants (ASVs) at 100% identity. Taxonomic assignments were performed using the SILVA v.138 database (Quast et al., 2013).

### Microbiota phenotypes

Microbiota phenotypes consisted of taxonomic composition and diversity metrics. Microbial abundance was determined by aggregating the non-rarefied sequence data at different taxonomic levels, ranging from genus to phylum, and normalizing them using the centered log-ratio (CLR). To ensure robustness and relevance in the analysis, only taxa that showed a relative abundance greater than 0.01% in at least one sample and were present in at least 50% of samples were considered. These selected taxa were subsequently included in the analysis of microbial phenotypes.

To mitigate the effect of sequence depth bias while estimating diversity indices, each sample was rarefied to 6,000 reads using the phyloseq (v1.40.0) and vegan (v2.6-2) R packages. The α-diversity indices - Shannon, Simpson, and Chao1 - were calculated using the rarefied dataset at both the ASV and genus levels. Furthermore, β-diversity was assessed using the Bray-Curtis dissimilarity matrix generated with the vegan R package (v2.6-2). Non-metric Multi-dimensional Scaling (NMDS) was performed on the Bray-Curtis dissimilarity matrix, and the values from the first two axes (NMDS1 and NMDS2) were used as β-diversity phenotypes.

### Estimation of variance components and genetic parameters

Microbiota phenotypic data were first analyzed using univariate animal models. Variance components were estimated using the AI-REML algorithm implemented in the Wombat software (Meyer, 2007). The linear animal model employed for this analysis was detailed as follows:

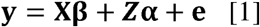

In the model, the vector **y** represented various microbiota phenotypes, CLR-transformed abundance of selected taxa, α-diversity indices (i.e., Shannon, Simpson, and Chao1), and, for β-diversity, values from the first two axes of NMDS (NMDS1 and NMDS2). The model also included a vector (**e)** for the random additive genetic effect, distributed as N (0**, A***σ_a_^2^*), and vector (e) for random residual effects, distributed as N (0, **I**σ_e_^2^). The matrices **x** and **z** served as the incidence matrices, and **A** represented the pedigree relationship matrix (four generations and 2,959 animals). **I** denoted the identity matrix. The parameters σ*_a_*^2^ and σ*_e_*^2^ were the additive genetic and residual variances, respectively. The **β** vector encompassed the fixed effects of several factors that were found significantly associated with the phenotypes, using analysis of variance (ANOVA) from the car package in R (v3.1-0) and permutational multivariate analysis of variance (PERMANOVA) from the vegan R package (v2.6-2). These factors included herd (18 herds), parity (five parities: 1^st^, 2^nd^, 3^rd^, 4^th^, and ≥ 5^th^), sampling season (three seasons), laboratory batch (four groups), and the time interval from calving to sampling (nine groups: 20-day intervals from 20 to 425 post-calving). Additionally, the number of sequence reads was included as a fixed effect in the model for taxa abundance, as this phenotype was derived from the non-rarefied dataset. The heritability (h^2^) was computed using additive genetic variance (σ*_a_*^2^) and residual variance (σ*_e_*^2^) according to the following formula.

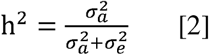

Covariances were estimated using bivariate models, in which both genetic and residual variances were represented as 2 × 2 matrices. We assessed genetic correlations among microbial taxa at all taxonomic levels, as well as between microbiota phenotypes and host traits related to fertility, production, and health. The additive genetic correlation (r*_α_*) was calculated in the bivariate models using the following formula:

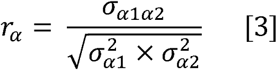

with *σ_α_*_1*α*2_, the additive covariance between traits one and two, and σ_α1_^2^ and σ_α2_^2^, the additive variance of traits one and two, respectively. Heritability and genetic correlation estimates were considered significant at *p* ≤ 0.05. *P*-values were calculated by comparing the test statistics to the Student’s t-distribution using the pt() function in R (R Core Team, 2022). To correct for multiple testing, we applied a 10% false discovery rate (FDR) threshold.

### Whole genome-wide association studies (GWAS) on heritable taxa and diversity indices

To identify genomic regions associated with microbiota phenotypes, GWAS were performed only on taxa and diversity indices with heritability estimates significantly different from zero. The 50K SNP genotypes of the 681 Holstein cows were imputed to whole-genome sequences (WGS) using a two-step approach to maximize imputation accuracy (Van Binsbergen et al., 2014). The imputation procedure has been described in detail by Sanchez et al. (2019). First, genotypes were imputed from 50K to 777K high-density (HD) SNPs using FImpute (Sargolzaei et al., 2014), and subsequently from HD SNPs to the whole-genome sequence (WGS) level using Minimac (Howie et al., 2012).

Imputation from the 50K to HD SNP level was carried out using a within-breed reference set of 776 Holstein bulls, genotyped with the Illumina BovineHD BeadChip (Hozé et al., 2013). WGS variants were imputed from HD SNP genotypes using WGS data from 3,414 *Bos taurus* animals, sourced from the ninth run of the 1000 Bull Genomes Project (Bouwman and Veerkamp, 2014; Brøndum et al., 2014). The imputation accuracy from HD SNPs to WGS was evaluated using the coefficient of determination (R^2^) calculated with Minimac. For further association analyses, only variants with an R^2^ value higher than 20% and a minor allele frequency (MAF) above 1% were retained. The number of variants (SNPs and InDels) ranged from 13,253,536 to 13,789,622 across the traits analyzed.

Single-trait association analyses were performed between the variants and the heritable taxa and diversity indices. Associations were tested using GCTA software (version 1.24) with the --*mlma* option, applying a mixed linear model (Yang et al., 2011).

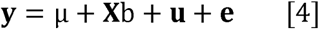

Where **y** represented the vector of adjusted CLR abundance of heritable taxa and diversity phenotypes, μ denoted the overall mean, and b represented the additive fixed effect of the variant under consideration. **x** was the matrix containing predicted allele dosage, ranging from 0 to 2. The vector **u** denoted the random polygenic effect, assumed to follow N (0, **G**σ*_u_*^2^), where **G** was the genomic relationship matrix (GRM) calculated using 50k genotypes and σ*_u_*^2^ was the polygenic variance estimated in the null model (Y = μ+ **u** + **e**). The vector **e** denoted random residual effects as assumed to follow N (0, **I**σ*_e_*^2^), where **I** was the identity matrix, and σ*_e_*^2^ was the residual variance. The genome-wide significance threshold was set at a nominal *P* value of 5×10^-8^ (−log_10_(*P*) = 7.3), as recommended in Sahana et al. (2023). This value corresponds to a Bonferroni correction based on one million independent tests and is relatively stringent given the long-range linkage disequilibrium patterns observed in cattle (Makanjuola et al., 2020; Gautason et al., 2021). Quantitative trait locus (QTL) confidence intervals were defined by the most distant variants located within the upper third of each QTL peak (Sanchez et al., 2017; Brulin et al., 2025). The percentage of phenotypic variance explained by each QTL for each trait was estimated as follows (Sanchez et al., 2019):

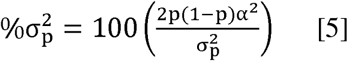

In this equation, σ_p_^2^ represents the phenotypic variance of the trait while α and p denote the estimated variant effect and frequency, respectively. Functional annotation was done using Ensembl (ver. Genes-112) (https://www.ensembl.org/Bos_taurus/biomart/).

## Results

### Characterization of vaginal microbiota phenotypes

A total of 173 taxa at various taxonomic levels were selected for microbiota phenotypes. These comprised nine phyla, 14 classes, 38 orders, 48 families, and 64 genera, each with a relative abundance > 0.01% in at least one sample and a prevalence > 50%. This filtering strategy ensured that only robust, biologically relevant taxa were included in downstream genetic analyses.

### Heritability of vaginal microbiota taxa and diversity indices

Heritability estimates across taxonomic levels ranged from 0.09 to 0.39 (|h² ± SE| > 0), as illustrated in Figure 1. These included five genera, three families, eight orders, three classes, and one phylum. Nominal statistical significance (*p* ≤ 0.05) was detected for one genus, one family, and three orders; however, none of these associations remained significant after false discovery rate (FDR) correction (FDR>0.10).

**Figure 1:**
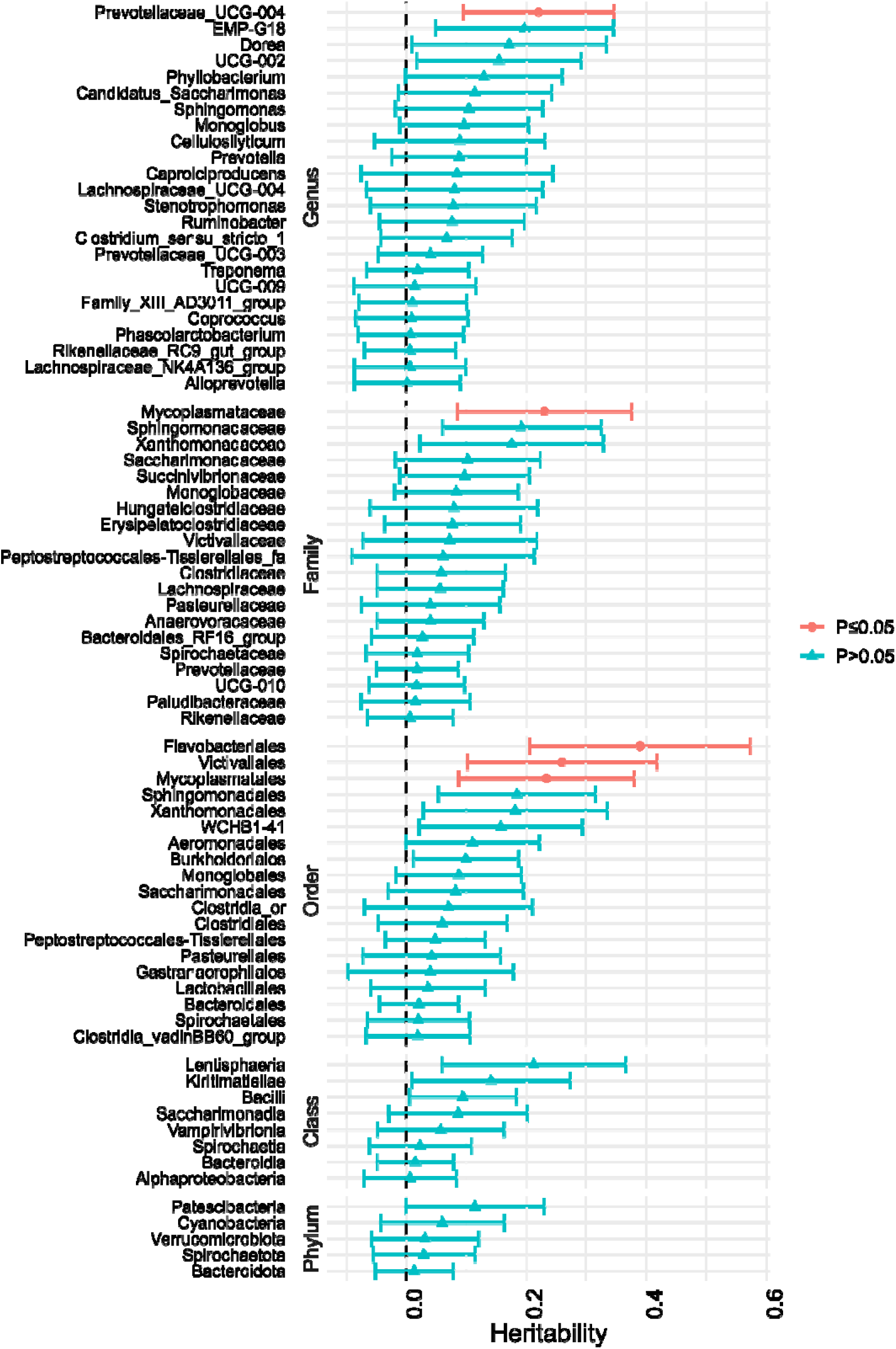
Heritability estimates of vaginal microbiota taxa across different taxonomic levels from genus to phylum. The vertical bars represent heritability, with only estimates for which the absolute value of h² ± SE exceeds zero (|h² ± SE| > 0) displayed. Red:

At the genus level, *Prevotellaceae_UCG-004* showed a detectable heritability (h^2^ = 0.22, SE = 0.12, *p* = 0.04). At the family level, *Mycoplasmataceae* was the only family with evidence of genetic influence (h^2^ = 0.23, SE = 0.14, *p* = 0.05). At the order level, Flavobacteriales (h^2^ = 0.39, SE = 0.18, *p* = 0.01), Victivallales (h^2^ = 0.26, SE = 0.16, *p* = 0.05), and Mycoplasmatales (h^2^ = 0.26, SE = 0.16, *p* = 0.05) exhibited notable heritability estimates.

At higher taxonomic levels, although no estimates reached nominal significance, several taxa presented non-zero heritability. For example, Lentisphaeria displayed the highest estimate among classes (h^2^ = 0.21, SE = 0.15, *p* = 0.08), while the most heritable phylum, Patescibacteria, presented a lower estimate (h^2^ = 0.11, SE = 0.11, *p* = 0.16). Additionally, the Chao1 diversity index (ASV level) had a low and non-significant heritability estimate (h² = 0.06, SE = 0.05). Other diversity metrics (Shannon, Simpson, NMDS axes) also yielded low and non-significant heritability values.

### Genetic and phenotypic correlations between microbiota phenotypes

Genetic and phenotypic correlations among heritable taxa are presented in Figure 2. In total, 59 non-null genetic correlations were estimated among genera, 30 among families, 33 among orders, three among classes, and two among phyla (|r_α_ ± SE| > 0). Nominal significance (p ≤ 0.05) was observed for 16 genus-level genetic, three class-level, and two order-level correlations, and one diversity-trait indice correlations. After applying multiple testing correction (FDR 10%), seven genus-level genetic correlations remained significant.

**Figure 2:**
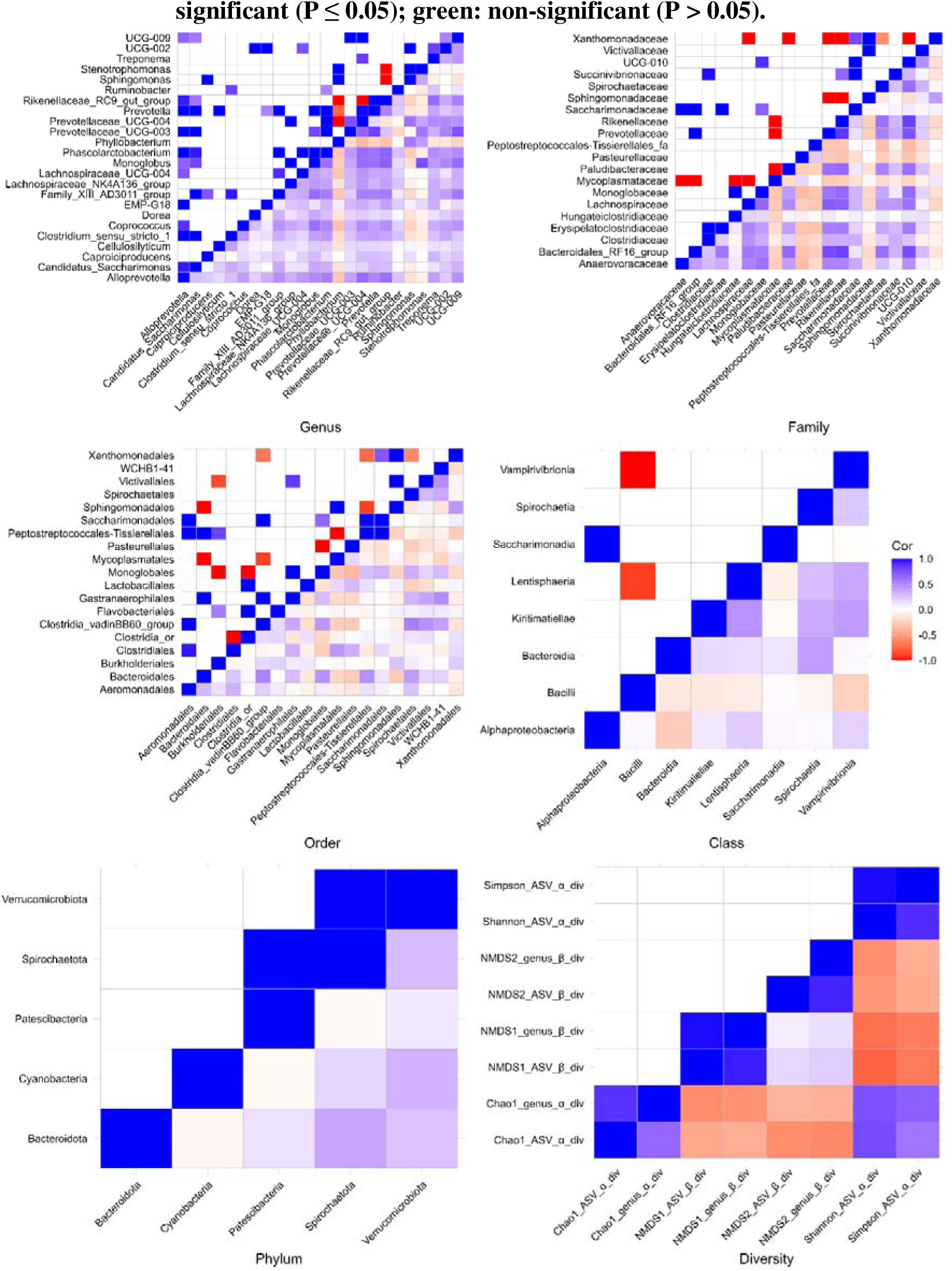
Genetic and phenotypic correlations between microbiota phenotypes (i.e., abundances and diversity indices). The upper left triangle indicates genetic correlation, and the lower right triangle indicates phenotypic correlation. The value of the correlations is represented by a color scale from blue (r = +1) to red (r = −1).

All 16 nominally significant genus-level genetic correlations were positive, ranging from 0.74 to nearly 1.00, with SEs ranging from 0.13 to 0.47. Notably, *Prevotellaceae_UCG-004* was involved in six of the seven FDR-significant genus-level correlations, showing strong positive genetic association with *Alloprevotella* (r ≈ 1.00, SE = 0.22, *p* = 4.3×10^-6^, *q* = 4.3×10^-4^), *Candidatus_Saccharimonas* (r ≈ 1.00, SE = 0.26, *p*= 1.4×10^-4^, *q* = 0.008), *Monoglobus* (r ≈ 1.00, SE = 0.14, *p* = 3.8×10^-13^, *q* = 1.5×10^-10^), *Family_XIII_AD3011_group* (r ≈ 1.00, SE = 0.23, *p* = 1.2×10^-5^, *q* = 7.7×10^-4^), *Phascolarctobacterium* (r ≈ 1.00, SE = 0.29, *p* = 6.2×10^-2^, *q* = 0.024), and *Rikenellaceae_RC9_gut_group* (r ≈ 1.00, SE = 0.14, *p* = 4.1×10^-11^, *q* = 5.6×10^-9^), respectively.

At the family level, two notable positive genetic correlations were observed between *Clostridiaceae* and *Succinivibrionaceae* (r_α_ = 0.98, SE = 0.42, *p* = 0.021) and between *Monoglobaceae* and *UCG-010* (r_α_ = 0.89, SE = 0.35, *p* = 0.012). Conversely, *Mycoplasmataceae* showed a strong negative genetic correlation with *Paludibacteraceae* (r_α_ ≈ −1.00, SE = 0.45, *p* = 0.027).

At the order level, Aeromonadales exhibited a high positive genetic correlation with Clostridiales (r_α_ = 0.96, SE = 0.37, *p* = 0.011), while Gastranaerophilales was positively correlated with Victivallales (r_α_ = 0.85, SE = 0.41, *p* = 0.031).

At higher taxonomic ranks, e.g., class and phylum levels, both strong positive and negative correlations were detected; however, none remained significant after FDR correction. At the class level, Alphaproteobacteria was positively correlated with Saccharimonadia (r_α_ ≈ 1.00, SE = 0.72), whereas Bacilli showed a strong negative correlation with Lentispheria (r_α_ = −0.99, SE = 0.69) and Vampirivibrionia (r_α_ ≈ −1.00, SE = 0.49). Among the phyla, Patescibacteria exhibited a positive genetic correlation with Spirochaetota (r_α_ ≈ 1.00, SE = 0.92), and Spirochaetota was also positively correlated with Verrucomicrobiota (r_α_ = 0.99, SE = 0.88). Regarding diversity phenotypes, a strong positive genetic correlation was identified between NMDS1_ASV and NMDS1_genus (r_α_ = 0.98, SE = 0.39, *p* = 0.013).

Phenotypic correlations across taxonomic levels were also assessed and were frequently significant (195 genera, 132 families, 71 orders, 16 classes, and seven phyla) even after FDR correction. These ranged from −0.36 to 0.81 for genus, −0.36 to 0.86 for family, −0.37 to 56 for order, −0.27 to 0.47 for class, 0.09 to 0.38 for phylum, and −0.76 to 0.94 for diversity phenotypes. Details of the phenotypic correlations with significant values are presented in Supplementary file 1.

### Genetic correlations between microbiota traits and host phenotypes

Genetic correlations between host phenotypes and heritable microbiota traits are shown in Figure 3. Non-zero genetic correlations (|r_α_ ± SE| > 0) were found in 18 genera, 14 families, 19 orders, 14 classes, five phyla, and 17 diversity-related phenotypes, each associated with at least one trait related to fertility, production, and health. Nominally significant associations (*p* ≤ 0.05) were observed between two genera, two families, one order, two classes, and four diversity traits showed a strong statistical association; however, none remained after FDR correction.

**Figure 3:**
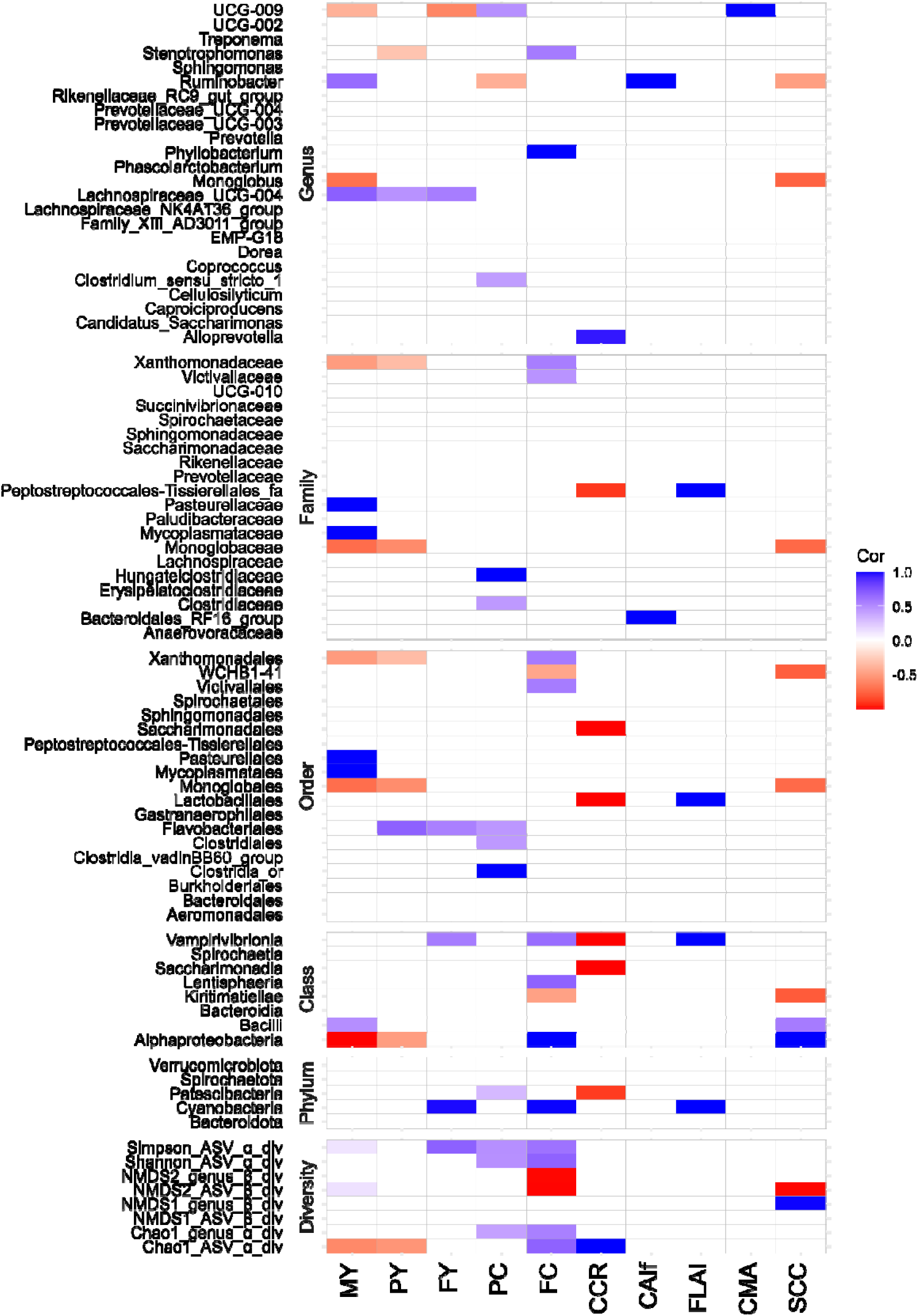
Genetic correlations between microbiota traits and host phenotypes. The value of the correlations is represented by a color scale from blue (r = +1) to red (r = −1).

At the genus level, positive genetic correlations were found between Lachnospiraceae_UCG-004 and MY (r_α_ = 0.69, SE = 0.36, *p* = 0.05), and between Stenotrophomonas and FC (r_α_ = 0.57, SE = 0.23, *p* = 0.01). At the family level, *Peptostreptococcales-Tissierellales_fa* was negatively corrected with CCR (r_α_ = −0.92, SE = 0.44, *p* = 0.04), while *Xanthomonadaceae* showed a positive correlation with FC (r_α_ = 0.56, SE = 0.26, *p* = 0.03). Similarly, at the order level, Xanthomonadales was positively correlated with FC (r_α_ = 0.57, SE = 0.26, *p* = 0.03). At the class level, Alphaproteobacteria was negatively correlated with MY (r_α_ ≈ −1.00, SE = 0.45, *p* = 0.03) and positively correlated with SCC (r_α_ ≈ 1.00, SE = 0.48, *p* = 0.04). Regarding diversity traits, FC was positively genetically correlated with four indices: chao1_ASV (r_α_ = 0.68, SE = 0.29, *p* = 0.01), NMDS2_ASV (r_α_ ≈1.00, SE = 0.51, *p* = 0.03), Shannon_ASV (r_α_ = 0.68, SE = 0.38, *p* = 0.04), and Simpson_ASV (r_α_ = 0.61, SE = 0.35, *p* = 0.04).

### Whole-genome GWAS analyses for vaginal microbiota

Whole-genome GWAS analyses performed with the 50K SNP array identified significant associations between genetic variants and heritable vaginal taxa and diversity phenotypes (|h² ± SE| > 0). Significant signals were detected for two genera (*Alloprevotella* and *Prevotellaceae_UCG-004*), two families (*Bacteroidales_RF16_group* and *UCG-010*), two orders (Bacteroidales and Xanthomonadales), one class (Bacteroidia), and one diversity trait (Simpson_ASV) (Figures 4 and 5).

**Figure 4:**
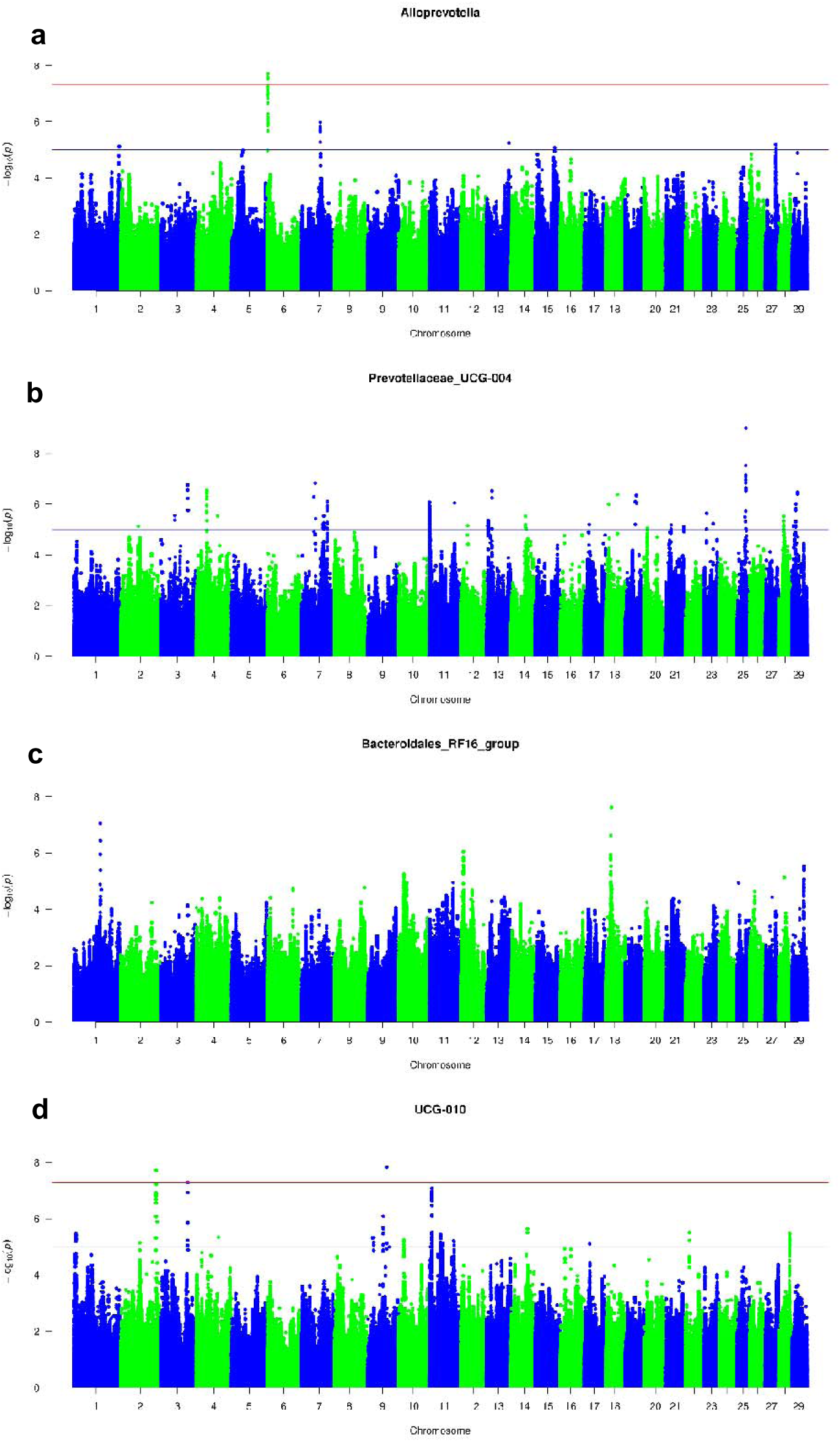
Manhattan plots showing −log_10_(*P*) values plotted against the positions of variants on cattle autosomes for the abundances of: Genus: (a) *Alloprevotella* (b) *Prevotellaceae_UCG-004*, Family: (c) *Bacteroidales_FR16_group* (d) *UCG-010*. The red line indicates the significant threshold level (−log_10_(*P*) = 7.3), and the blue line indicates the suggestive threshold (−log_10_(*P*) = 5.0).

**Figure 5:**
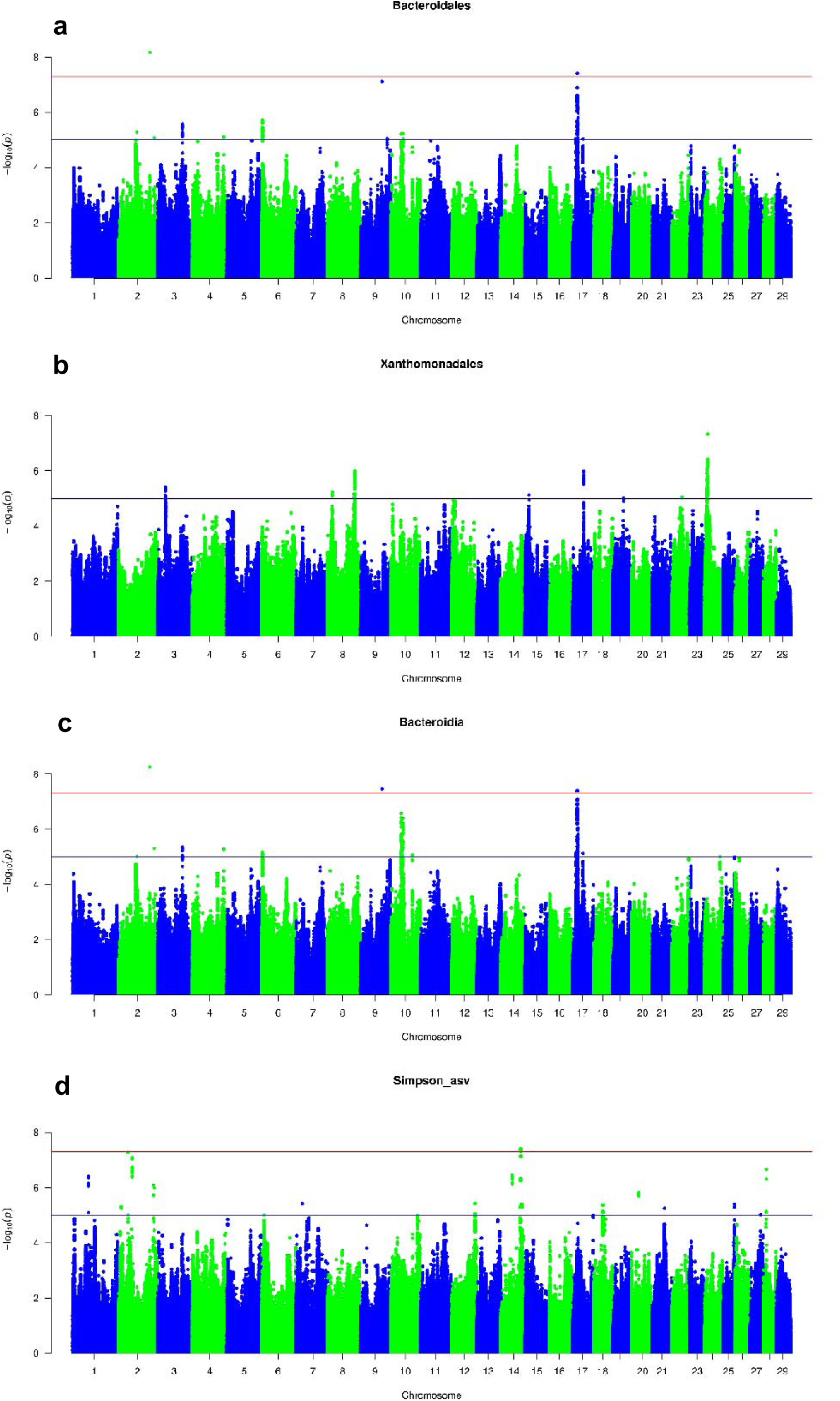
Manhattan plots showing −log_10_(*P*) values plotted against the positions of variants on cattle autosomes for the traits. Order: (a) Bacteroidales (b) Xanthomonadales, Class: (c) Bacteroidia, Diversity: (d) Simpson_asv. The red line indicates the significant threshold level (−log_10_(*P*) = 7.3), and the blue line indicates the suggestive threshold (−log_10_(*P*) =5.0).

At the genus level, *Alloprevotella* showed a significant QTL on *Bos taurus* (BTA) autosome 6 with −log_10_(*P*) = 7.3. Within the 2 Mbp region defining this QTL, 24 variants were located in the upper third of the peak. The lead candidate variant, rs2619566 (−log_10_(*P*) = 7.71), was located at 2,619,566 bp within the *NAFI* gene region and explained 6.6% of the phenotypic variance (Table 2). *Prevotellaceae_UCG-004* exhibited a significant peak on BTA25. Within this QTL region, five variants were identified, with rs29029332 emerging as the lead candidate variant (−log_10_(*P*) = 9.00), located in the *CALN1* gene (Table 2).

**Table 2:** QTL region and best candidate variant identification by GWAS (a) QTL region (b) Candidate variant.

| Trait | Rank | BTA | From (bp) | To (bp) | Variants | Gene |
| --- | --- | --- | --- | --- | --- | --- |
| *Alloprevotella* | Genus | 6 | 1915612 | 2724965 | 129 | 5 |
| *Prevotellaceae_UCG-004* | Genus | 25 | 28411547 | 29136217 | 9 | 1 |
| *Bacteroidales_RF16_group* | Family | 18 | 17517825 | 19488872 | 11 | 11 |
| *UCG-010* | Family | 2 | 120737853 | 121133689 | 14 | 11 |
|  | Family | 9 | 64777489 | 64816319 | 2 | - |
| Bacteroidales | Order | 17 | 15899853 | 16909624 | 39 | 3 |
| Xanthomonadales | Order | 24 | 8499084 | 10833625 | 70 | 4 |
| Bacteroida | Class | 17 | 15184475 | 17560658 | 47 | 10 |
| Simpson_asv | α-diversity | 14 | 68789014 | 69289184 | 10 | 4 |

**b. Top candidate variant**
| Trait | BTA | Variant Id | $-\log_{10}(P)$ | Freq | Effect | SE | % Phenotypic variance | Gene | Functional annotation |
| --- | --- | --- | --- | --- | --- | --- | --- | --- | --- |
| *Alloprevotella* | 6 | rs2619566 | 7.71 | 0.98 | 15.77 | 2.18 | 6.57 | *NAF1* | Upstream |
| *Prevotellaceae_UCG-004* | 25 | rs29029332 | 9.0 | 0.01 | -31.13 | 5.10 | 34.23 | *CALN1* | Intron |
| *Bacteroidales_RF16_group* | 18 | rs1948872 | 7.61 | 0.23 | -3.55 | 0.63 | 5.58 | - | Intergenic |
| *UCG-010* | 2 | rs121114033 | 7.71 | 0.01 | -28.13 | 5.01 | 14.34 | - | Intergenic |
|  | 9 | rs64816319 | 7.83 | 0.98 | 22.92 | 4.04 | 18.85 | - | Intergenic |
| Bacteroidales | 17 | rs16104650 | 7.41 | 0.10 | -5.10 | 0.92 | 5.42 | *IL15* | Intron |
| Xanthomonadales | 24 | rs10496315 | 7.32 | 0.03 | -12.64 | 2.31 | 10.43 | - | Intergenic |
| Bacteroida | 17 | rs16104650 | 7.39 | 0.10 | -4.99 | 0.91 | 5.38 | *IL15* | Intergenic |
| Simpson_asv | 14 | rs69272114 | 7.41 | 0.01 | -6.45 | 1.17 | 52.46 | *NDUFAF6* | Intron |

\* BTA= *Bos taurus* autosome; bp= Base pair; Freq= Frequency; SE= Standard error

At the family level, the *Bacteroidales_RF16_group* showed a significant QTL on BTA 18, with rs1948872 being the lead candidate variant (−log_10_(*P*) = 7.61). Besides, two significant QTL regions were found associated with UCG-010 abundance on BTA2 and BTA9, with rs121114033 and rs64816319 emerging as lead variants (−log_10_(*P*) = 7.71 and 7.81, respectively) (Table 2).

At the order level, both Bacteroidales and Xanthomonadales presented a significant association with host genomic regions. For Xanthomonadales, a QTL was identified on BTA24. Among the 70 variants within the QTL confidence interval, rs10496315 was identified as the lead variant (−□log_10_(*P*)□=□7.32), explaining 10.4% of phenotypic variance. Bacteroidales showed two significant QTL on BTA2 and 17. On BTA17, among the 39 variants in the QTL confidence interval, rs16104650 was the lead candidate (−log_10_(*P*) = 7.41) in the *IL15* gene region. At the class level, Bacteroidia also displayed significant associations with BTA2, BTA9, and BTA17. Notably, the associated order, Bacteroidales, showed overlapping genomic regions on the same chromosomes. In particular, a similar confidence interval for BTA17 was observed, in which rs16104650 again emerged as the lead candidate variant (−log10(P) = 7.39), reinforcing the relevance of the *IL15* gene at both taxonomic levels.

Finally, the only diversity index tested, i.e., Simpson_asv, showed a significant QTL on BTA14. The QTL confidence interval comprised 10 variants, with rs69272114 within the *NDUFAF6* gene, identified as the lead variant (−log_10_(*P*) = 7.41) (Table 2).

## Discussion

Our objectives were to gain a better understanding of the genetic and genomic relationships between the cow and one of its less-studied microbiota, the vaginal microbiota. Although a moderate number of cows were sampled, this study identified low to moderate heritabilities for some vaginal microbiota traits and detected genetic correlations with host traits in Holstein cows. To the best of our knowledge, this study represents the first investigation of the relationship between host genetics and vaginal microbiota in cows. We also identified specific genes and genomic variants associated with microbiota traits through genome-wide association studies (GWAS). Our findings suggest that host genetics contributes to vaginal microbiota composition and diversity, as well as to genetic correlations with fertility, production, and health-related phenotypes, with potential applications in the bovine industry.

### Host genetics influences on vaginal microbiota composition

The heritability (h²) estimates ranged from 0.09 to 0.39, indicating that genetic factors could account for a limited to moderate proportion of the variance in microbial abundance.

At the genus level, *Prevotellaceae_UCG-004* exhibited the highest heritability (0.22 ± 0.12), indicating a moderate genetic influence. Other genera with detectable heritability included *EMP-G18* (0.19 ± 0.14), *Dorea* (0.17 ± 0.16), and *UCG-002* (0.15 ± 0.13). Notably, these genera belong to the Firmicutes and Bacteroidetes phyla, which have also been reported to include heritable vaginal microbial groups in sows. For instance, Sanglard et al. (2020) reported that the most prevalent Operational Taxonomic Units (OTUs) associated with the Firmicutes, Proteobacteria, and Bacteroidetes phyla demonstrated moderate to high heritability (h^2^ > 0.20). These cross-species similarities suggest that host genetic factors may influence microbial groups within dominant phyla, independent of host species. In contrast, extensive research has examined the relationship between host genetics and vaginal microbiota in humans (Si et al., 2017; Wright et al., 2021). A study using the Korean twin-families dataset identified *Prevotella* (h^2^ = 0.72) as the vaginal microbiota genus with the highest heritability, followed by *Lactobacillus iners* (h^2^ = 0.41) and *L. crispatus* (h^2^ = 0.36) (Si et al., 2017). Another study, conducted by Wright et al. (2021), found a strong genetic influence on *L. crispatus* presence among women of European descent, with a heritability estimate of 0.34. Collectively, these findings indicate that host genetic factors influence the composition of vaginal bacterial genera.

Consistent with previous analyses of the same cohort (Brulin et al., 2024a), which demonstrated detectable heritability for the genus *Prevotellaceae_UCG-004* in fecal microbiota (h^2^ = 0.12), the present study identified a higher heritability estimate for the same genus in vaginal samples (h^2^ = 0.22). This difference may result from stronger host genetic control in the vaginal niche, lower environmental variability relative to the gut, and niche-specific host–microbiota interactions. Additionally, statistical factors, such as sample size and differences in variance structure across sample types, may contribute to the observed variation in heritability estimates. The recurrence of this heritable taxon across multiple anatomical sites supports the hypothesis that host genetic factors play a role in shaping microbial community composition. However, because an overlapping taxon was detected in both fecal and vaginal samples, limited cross-site contamination during sampling cannot be entirely excluded and should be considered when interpreting these findings.

The heritability estimate was the highest for the Flavobacteriales order (0.38±0.18), suggesting a significant genetic contribution to its abundance. Lentisphaeria exhibited the highest heritability (0.21 ± 0.15) at the class level, while Kiritimatiellae had a slightly lower heritability (0.14 ± 0.13). These findings indicated that higher taxonomic levels were more strongly influenced by the host’s genetic makeup, which is essential for understanding broader trends in microbial inheritance and their effects on host health. The heritability values of rumen bacterial diversity indicators, such as α and β-diversities, ranged from 0.15 to 0.23 (Li et al., 2019). The heritability estimate for the Chao1 diversity index in our study was low (0.06 ± 0.05), indicating that genetic factors have a minimal influence on total microbial diversity at the ASV level. Alternative diversity metrics, such as Shannon, Simpson, and the NMDS axes derived from Bray-Curtis distances, did not exhibit substantial heritability, as also noted in the fecal microbiota (Brulin et al., 2024a). These findings suggest that non-genetic variables, such as environmental conditions and management approaches, primarily influence microbial diversity (Rothschild et al., 2018).

Multiple testing correction was applied, consistent with previous studies on rumen and fecal microbiota in cattle (Difford et al., 2018; Brulin et al., 2024a). No significant heritable taxa were observed in vaginal swabs of Holstein cows following FDR correction. The lack of significance is likely attributed to the relatively limited sample size, as FDR correction is recognized as conservative, particularly in microbiota datasets with numerous taxa and fewer animals (Difford et al., 2018; Brulin et al., 2024a). Furthermore, a substantial proportion of the variation in vaginal microbiota appears to be influenced by environmental factors rather than host genetics. Nevertheless, several taxa with low to moderate heritabilities were identified, indicating that host genetics may still contribute to the composition of vaginal microbial communities. Although these findings are preliminary, such heritable taxa may be of interest because vaginal microbiota can influence reproductive health, disease susceptibility, and overall host fitness (Adnane and Chapwanya, 2022; Amat et al., 2022; Ni et al., 2023). Future studies involving larger cohorts and multiple breeds are necessary to validate these results and to assess their potential relevance in breeding programs aimed at improving animal health and resilience.

### Genetic and phenotypic correlations between vaginal microbiota phenotypes and diversity

The study identified substantial genetic and phenotypic correlations among heritable microbial taxa across multiple taxonomic levels and diversity phenotypes. Pronounced genetic correlations between microbiota taxa (ranging from nearly −1 to +1) highlight the complex genetic architecture influencing the vaginal microbiota in cattle. Phenotypic correlations were of lower magnitude than genetic correlations, a trend also observed in the human vaginal microbiota (France et al., 2022).

Interestingly, most of the genera that showed a substantial positive correlation did not belong to the same family. *Prevotellaceae_UCG-004*, a member of the *Prevotellaceae* family, exhibited strong positive genetic correlations nearly to 1.00 with other genera, including *Candidatus_Saccharimonas* (*Saccharimonadaceae* family), *Monoglobus* (*Monoglobaceae* family), *Family_XIII_AD3011_group* (*Anaerovoracaceae* family), *Phascolarctobacterium* (*Acidaminococcaceae* family), and *Rikenellaceae_RC9_gut_group* (*Rikenellaceae* family). Conversely, *Phyllobacterium* from the *Rhizobiaceae* family showed negative genetic correlations with *Provetellaceae_UCG-004* (*Prevotellaceae* family). This indicated that these genera still displayed genetic correlations despite belonging to different families. The genetic relationships observed among various genera underscored that microbial community formation can be influenced by intricate ecological interactions that extend beyond mere taxonomic similarity (Ghoul and Mitri, 2016; Louca et al., 2018).

Similar findings were also noticed in humans, where genera clustered into broader community groups despite taxonomic distance, as reported in the vaginal community state type studies (Ravel et al., 2011; Ceccarani et al., 2019). Studies on microbial enterotypes indicate that selecting for one or two keystone taxa can restructure the entire microbial community, as demonstrated by Larzul et al. (2024), who reported that selecting for specific enterotypes significantly altered gut microbiota composition in pigs. These findings suggest that similar principles may govern the bovine vaginal microbiota. Correlations and associations across taxonomic levels indicate that targeting specific taxa can influence overall community composition and diversity.

### Genetic correlations between microbiota traits and host phenotypes

The findings of this study underscore the complexity of these associations, as notable genetic correlations were observed across taxonomic levels and diversity phenotypes. To date, no studies have examined the genetic relationships between vaginal microbiota and host traits in dairy cows. However, evidence from human studies suggests that host genetics partially influence vaginal microbiota composition and that specific taxa exhibit genetic correlations with reproductive and infection-related phenotypes. Twin-based analyses showed that several vaginal taxa, including *Prevotella* dominated communities, were heritable and associated with metabolic and infection susceptibility traits (Si et al., 2017). Additionally, genomic studies demonstrated that variation in immune-related pathways contributes to differences in vaginal microbiome structure (Mehta et al., 2020). Furthermore, research on dairy cattle provides information on the genetic links between the rumen and fecal microbiota and the host’s phenotypic traits, including feed efficiency, milk production, and methane production (Jami et al., 2014; Difford et al., 2018; Li et al., 2019; Brulin et al., 2024a). Collectively, these studies suggest that specific microbial groups, rather than the overall microbiota composition, may play a critical role in influencing host performance.

For fertility traits, the genus *Alloprevotella* demonstrated a robust positive correlation with CCR, suggesting a potential role in reproductive success. While direct evidence connecting *Alloprevotella* to fertility in cattle remains limited, studies in other hosts suggested plausible mechanisms. In humans, elevated *Alloprevotella* levels have been observed in certain reproductive disorders, such as endometriosis, which further associates this genus with reproductive tract health (Miyashira et al., 2022). Collectively, these findings support the hypothesis that variation in *Alloprevotella* abundance may influence fertility; however, targeted functional and longitudinal studies in bovine vaginal microbiota are required to establish causality. At the family level, *Bacteroidales_RF16_group* was positively correlated with CAIf, indicating that this family might contribute to more efficient reproductive processes. A study investigating the genetic relationships between fecal microbiota and reproductive parameters in Holstein cows found consistent results. Specifically, the relative abundance of the genus *Alloprevotella* in the microbiota was positively correlated with CCR, whereas the abundance of the genus Bacteroidales_RF16_group was positively correlated with CAIf (Brulin et al., 2024a). The consistent findings across two different bacterial communities in cows emphasize the potential influence of these communities on cattle’s reproductive traits.

Additionally, the study revealed substantial genetic correlations between microbiota and dairy production traits. Correlations with production traits ranged from notable positive to moderate negative (0.69 to −0.41) across different taxonomic levels, reflecting the complex interplay between the microbiota and production efficiency. The *Stenotrophomonas* genus and the *Xanthomonadaceae* family were positively correlated with fat content, and the *Lachnospiraceae_UCG-004* genus was positively correlated with milk yield. However, unlike fertility traits, some taxa did not exhibit consistent associations across body sites. Although we found no association between the *Prevotellaceae* genus and production traits in vaginal samples, previous studies on rumen and fecal microbiota have reported negative genetic associations between this genus and production traits, such as milk and fat yields (Zhang et al., 2022; Brulin et al., 2024a). These findings underscore the importance of considering host location and prior research when interpreting microbiota effects.

Notable negative correlations were identified between the *Monoglobus* genus and its corresponding family (*Monoglobaceae*) and order (Monoglobales), as well as between the *Monoglobus* genus and somatic cell score (SCS), an indicator of mastitis and overall udder health. This negative association suggests that these taxa may help maintain udder health. Conversely, the *UCG-009* genus exhibited a positive correlation with clinical mastitis, indicating a potential role in increased disease susceptibility. Evidence from other species further substantiates the association between microbiota composition and health. Human studies have demonstrated links between specific bacterial taxa, such as *Prevotella*, and conditions including bacterial vaginosis and obesity (Si et al., 2017). *Lactobacillus spp.* are recognized for protecting the vaginal environment by inhibiting pathogenic bacteria (Boskey et al., 2001; Voravuthikunchai et al., 2006). Similarly, research in commercial gilts has shown that vaginal microbial composition influences immunological responses (Sanglard et al., 2020). Additionally, fecal microbiota analyses in dairy cows have revealed genetic associations between microbial taxa and immune-related traits, such as inflammatory markers and antibody levels (Brulin et al., 2025), reinforcing the significance of host–microbiome interactions in shaping immune function. Collectively, these findings indicate that vaginal microbiota composition is associated with multiple health aspects in both animals and humans. The results suggest that targeted modulation of the microbiome may enhance fertility, production, and health traits in cattle. For example, promoting beneficial microbes, such as *Alloprevotella* and *Peptostreptococcales-Tissierellales_fa*, may improve reproductive efficiency, while reducing detrimental taxa, such as *UCG-009*, could enhance disease resistance.

### Whole-genome GWAS studies of microbiota traits

This study highlights specific genomic regions within the host genome that are associated with features of the vaginal microbiota. These results offer new insights into the genomic mechanisms underlying the heritability and diversity of microbiota compositions. The following section focuses on four major signals identified on BTA6, BTA25, BTA17, and BTA14.

A QTL associated with the *Alloprevotella* genus was identified on BTA6, with top genetic variants located within the *NAF1* gene area. The *NAF1* gene is essential for ribosome synthesis (Fu and Collins, 2007; Egan and Collins, 2012) and is expressed in vaginal tissue according to the Cattle Genotype-Tissue Expression (cGTEx) portal (https://cgtex.roslin.ed.ac.uk/search/). Furthermore, *Alloprevotella* demonstrated a genetic correlation with fertility traits in this study, suggesting that host genomic regulation of this taxon may influence reproductive performance.

Furthermore, the most heritable genus, *Prevotellaceae_UCG-004* (h^2^=0.22), showed a significant association with genomic variants on BTA25, with prominent signals within the *CALN1* gene. *CALN1* encodes a calcium-binding protein that is primarily expressed in the brain (Kobuke et al., 2018). In cattle, *CALN1* expression occurs in the uterus (cGTEx portal), whereas in humans, it is reported in the vagina (human GTEx database). Additional functional studies are required to elucidate the mechanistic relationship between host *CALN1* variation and *Prevotellaceae_UCG-004* abundance. The leading variant accounted for 34.2% of phenotypic variance. However, this portion should be interpreted with caution, as effect estimates and variance explained may be inflated in studies with small sample sizes or variants with low minor allele frequency (MAF). Replication in larger and independent cohorts is essential to confirm the magnitude and robustness of this association.

On BTA17, two signals were identified that were associated with the Bacteroidales order and the Bacteroida class, both mapping to the *Interleukin 15* (*IL15*) locus. *IL15* is a key cytokine involved in mucosal and reproductive tract immunity; it regulates natural killer (NK) cells, memory T cells, and epithelial barrier function, and is essential for immune homeostasis in the uterus and placenta (Gordon, 2021). Evidence from animal models further supports the role of *IL15* in shaping the microbiota. Specifically, overexpression of *IL15* in mice alters gut microbial composition, leading a decreased butyrate-producing bacteria and increased susceptibility to inflammatory challenges (Meisel et al., 2017). Collectively, these findings suggest that host genetic variation at *IL15* may influence vaginal microbiota composition, potentially through the immune environment; however, direct evidence in cattle remains to be established.

A significant association was identified between Simpson_ASV and genetic variants on BTA14. The lead variant was located within the *NDUFAF* gene, which is expressed in bovine vagina tissue. *NDUFAF6* encodes a mitochondrial complex I assembly factor, suggesting a role in mitochondrial function and energy metabolism (Lemire, 2017). Currently, no direct evidence links *CALN1*, *NAF1*, or *NDUFAF6* to the regulation of the bovine vaginal microbiota. In contrast, human studies have identified genomic regions associated with specific features of the vaginal microbiome, such as *L. crispatus* and *L. iners*. Genes involved in innate immunity, including *TLR2*, *TIRAP*, and *MBL2*, have also been linked to these microbiota traits (Mehta et al., 2020). Evidence from cattle further demonstrates that host genetics influence gut and fecal microbial communities. For example, Brulin et al. (2025) reported genomic associations between fecal microbiota and immune-related pathways, and several rumen microbiota studies have identified heritable taxa and QTLs enriched for immune and metabolic functions (Li et al., 2019; Boggio et al., 2022; Wang et al., 2024). In pigs, QTLs associated with vaginal microbiota, including *Acinetobacter*, *Ruminococcaceae*, and *Clostridiales*, have been identified, with associated genes primarily related to immune system functions (Sanglard et al., 2020). Taken together, these findings suggest that immune and metabolic genes may contribute to shaping microbial community structure across multiple anatomical sites, although functional validation within the bovine vaginal tract remains to be demonstrated.

## Limitations and perspectives

Our study demonstrated a significant role of host genetics in regulating vaginal microbiota composition and identified substantial genetic correlations between microbiota traits and key host phenotypes. However, several limitations should be acknowledged. The relatively moderate sample size may have reduced statistical power and limited the precision of genetic parameter estimates. Future research should investigate additional breeds and environmental conditions to assess whether the genetic regulation of vaginal microbiota is conserved across populations or breed-specific. Furthermore, expanding phenotypic data collection, especially for health traits that are challenging to monitor routinely (such as reproductive tract infections and subclinical uterine disorders), would improve our understanding of the microbiota’s contribution to animal health. Advances in whole-genome metagenomics and host-microbiome multi-omics approaches will provide higher-resolution insights into microbial functions and their interactions with host biological pathways. Functional studies assessing gene expression, immune responses, and microbial activity are necessary to elucidate the mechanisms by which host genetic variants shape microbial communities. Incorporating microbiota data into breeding programs, either through microbiome-informed selection indexes or by directly including microbial traits in genetic evaluation models (Enciso et al., 2021; Álvaro et al., 2024), offers a promising strategy to improve fertility, production, and health outcomes. Investigating genetic correlations among vaginal, fecal, and ruminal microbiota will clarify whether host genetics exerts a systemic influence across body sites or whether site-specific genetic mechanisms shape distinct microbial communities.

## Conclusions

This study successfully identified microbial taxa under host genetic influence within the vaginal microbiota of dairy cattle, highlighting significant genetic correlations between these microbial communities and host phenotypes related to fertility, health, and production traits. Through comprehensive whole-genome GWAS, specific genetic variants associated with vaginal microbiota traits were identified, providing new insights into the genetic mechanisms shaping microbial composition and diversity. These findings have significant implications for cattle breeding strategies, suggesting that targeted selection for beneficial microbiota could improve overall health, fertility, and productivity in dairy cattle.

## Acknowledgements

The authors thank both the breeders and the technicians who collected the samples, as well as Gérard Heumez (Gènes Diffusion, Douai, France) for his assistance with data processing. The author also thanks Mekki Boussaha (Université Paris-Saclay, AgroParisTech, INRAE, GABI, Jouy-en-Josas, France) for his help and support in the imputation steps and the processing of the 1000 Bull Genome Project sequences. The authors gratefully acknowledge the French National Database (CTIG; Jouy-en-Josas, France) and Valogene (Paris, France) for providing data, and the contribution of the 1000 Bull Genomes Project. The authors additionally thank the INRAE MIGALE Bioinformatics Facility (MIGLE, INRAE, 2020) and the MIGALE Bioinformatics Facility (Jouy-en-Josas, France, doi: 10.15454/1.5572390655343293E12) for providing the computational resources needed for our analyses. GD Biotech expresses gratitude to the ANR-10-EQPX-07-01 LIGAN-PM-UMR8199 (Institut Pasteur de Lille, Lille, France) for its technical support regarding the MiSeq sequencing. AAMS extends my appreciation to the PRIAM masters and EMABG colleagues for their encouragement and contributions.

## Authors contribution

CA and SD conceptualized the study and designed the experiment. SMa and SMe conducted the whole lab work. AAMS performed the bioinformatic data processing and biostatistical analyses under the supervision of LB. AAMS, LB, JE, SD, PC, CA, and M-PS all participated in interpreting the results. AAMS drafted the manuscript. All authors conceptualized the manuscript, edited later drafts, read, and approved the final manuscript.

## Ethics

This study was conducted on dairy cattle reared for commercial purposes in compliance with the French Regulation (Code Rural et de la Pêche Maritime) and the European Council Directive 98/58/EC. The sampling of the animals was part of the routine animal manipulations carried out by duly authorized technicians from the Gènes Diffusion breeding company (Douai, France). Thus, under French law, no approval from an Institutional Animal Care and Use Committee or ethics committee was required. Additionally, the farmers involved in this study agreed to the use of their animals’ samples for research purposes.

## Nonstandard abbreviations used

ASV: amplicon sequence variant
CLR: centered-log ratio
CAIf: calving-to-fertilization interval via AI
FLAI: interval between the first and last AI
CCR: conception rate
MY: milk yield
PY: protein yield
FY: fat yield
PC: protein content
FC: fat content
CMA: clinical mastitis
FDR: false discovery rate
FDR 10%: FDR threshold of 10%
NMDS: nonmetric multidimensional scaling
YD: yield deviation
PERMANOVA: permutational multivariate analysis of variance
WGS: whole genome sequence
OTU: operational taxonomic unit
MAF: minor allele frequency
NK: natural killer
ILI5: interleukin 15

